# Human local adaptation of the TRPM8 cold receptor along a latitudinal cline

**DOI:** 10.1101/251033

**Authors:** Felix M. Key, Muslihudeen A. Abdul-Aziz, Roger Mundry, Benjamin M Peter, Aarthi Sekar, Mauro D’Amato, Megan Y. Dennis, Joshua M. Schmidt, Aida M. Andrés

**Affiliations:** Department of Evolutionary Genetics, Max Planck Institute for Evolutionary Anthropology, 04103 Leipzig, Germany; Max Planck Institute for Evolutionary Anthropology, 04103 Leipzig, Germany; Department of Human Genetics, University of Chicago, Chicago IL 60637, USA; Genome Center, MIND Institute, and Department of Biochemistry & Molecular Medicine, University of California, Davis, CA 95616, USA; BioDonostia Health Research Institute and IKERBASQUE, Basque Foundation for Science, San Sebastian, Spain

**Author notes:** Present Address: Department of Archaeogenetics, Max Planck Institute for the Science of Human History, 07745 Jena, Germany (Felix M. Key). Present address: Australian Centre for Ancient DNA, School of Biological Sciences and The Environment Institute, The University of Adelaide, Adelaide, SA, Australia. Present address: UCL Genetics Institute, Department of Genetics, Evolution and Environment, University College London, London, UK. **Correspondence** Felix M. Key and Aida M. Andrés.

## Abstract

Ambient temperature is a critical environmental factor for all living organisms. It was likely an important selective force as modern humans recently colonized temperate and cold Eurasian environments. Nevertheless, as of yet we have limited evidence of local adaptation to ambient temperature in populations from those environments. To shed light on this question, we exploit the fact that humans are a cosmopolitan species that inhabits territories under a wide range of temperatures. Focusing on cold perception – which is central to thermoregulation and survival in cold environments— we show evidence of recent local adaptation on *TRPM8.* This gene encodes for a cation channel that is, to date, the only temperature receptor known to mediate an endogenous response to moderate cold. The upstream variant rs10166942 shows extreme population differentiation, with frequencies that range from 5% in Nigeria to 88% in Finland (placing this SNP in the 0.02% tail of the F_ST_ empirical distribution). When all populations are jointly analysed, allele frequencies correlate with latitude and temperature beyond what can be explained by shared ancestry and population substructure. Using a Bayesian approach, we infer that the allele originated and evolved neutrally in Africa, while positive selection raised its frequency to different degrees in Eurasian populations, resulting in allele frequencies that follow a latitudinal cline. We infer strong positive selection, in agreement with ancient DNA showing high frequency of the allele in Europe 3,000 to 8,000 years ago. rs10166942 is important phenotypically because its ancestral allele is protective of migraine. This debilitating disorder varies in prevalence across human populations, with highest prevalence in individuals of European descent –precisely the population with the highest frequency of rs10166942 derived allele. We thus hypothesize that local adaptation on previously neutral standing variation may have contributed to the genetic differences that exist in the prevalence of migraine among human populations today.

**Author Summary:** Some human populations were likely under strong pressure to adapt biologically to cold climates during their colonization of non-African territories in the last 50,000 years. Such putative adaptations required genetic variation in genes that could mediate adaptive responses to cold. *TRPM8* is potentially one such gene, being the only known receptor for the sensation of moderate cold temperature. We show that a likely regulatory genetic variant nearby *TRPM8* has several signatures of positive selection rising its frequency in Eurasian populations during the last 25,000 years. While the genetic variant was and is rare in Africa, it is now common outside of Africa, with frequencies that strongly correlate with latitude and are highest in northern European populations. Interestingly, this same genetic variant has previously been strongly associated with migraine. This suggests that adaptation to cold has potentially contributed to the variation in migraine prevalence that exists among human groups today.

## Introduction

While our ancestors lived in Africa for millions of years, their successful colonization of colder environments outside of Africa is relatively recent, occurring during the last ~50,000 years. A number of novel genetic adaptations in populations that settled extreme polar environments are documented [1‐3]. This includes an allele in the gene *CPT1A*, which encodes a protein involved in the regulation of mitochondrial oxidation of fatty acids, in Northern Siberian populations [1, 2], and several alleles in genes involved in fatty acid metabolism in Greenlanders [3, 4]. These genetic changes likely represent adaptations to the highly specialized diets of these specific populations, which are rich in fatty acids. However, the putative adaptations to temperature and climate are largely unresolved.

Even in non-polar environments, temperatures range substantially across human habitats. For example, average annual temperature is 28°C in Nigeria (home to the Yoruba) and only 6°C in Finland, with differences most pronounced from December to February (29°C in in Nigeria and -4°C in Finland). These temperature differences illustrate the habitat changes experienced by early human groups as they migrated north. Local adaptation has significantly contributed to population differentiation that exists among human populations [5]. So it is reasonable to expect that besides genetic adaptations to selective factors that correlate with climate, such as diet [1‐3] and subsistence strategy [6], or pathogens [7] and their load [8], humans may harbour direct genetic adaptations to temperature and other climatic factors [6, 9].

Thermosensation (the sensation of innocuous environmental temperature) is crucial for thermoregulation (the process that maintains core body temperature) and is mediated by warm and cold receptor nerves that innervate the skin. At the molecular level, temperature sensation is due to the activation of transient receptor potential (TRP) ion channels. Among the few TRPs with clear thermoregulatory role (reviewed in [10]), only TRP cation channel subfamily M member 8 (TRPM8) is broadly agreed to play a central role in cold sensation and subsequent physiological thermoregulation [11‐17]. *TRPM8* is expressed in pain and temperature-sensitive neurons of the dorsal root ganglia [15], and at lower levels in other tissues such as prostate or liver [10, 18]. From approximately 15°C to 30°C the channel passes a mixed inward cationic current at cool to cold temperatures with strength inversely proportional to temperature. Interestingly, it is also activated by natural ligands such as menthol [17, 19] and is responsible for the local cooling sensation of mint-containing products [19]. Proof of its physiological role is that its deletion diminishes responses to cold [11‐13] including behavioural responses to innocuous cool, noxious cold, injury-evoked cold hypersensitivity and cooling-mediated analgesia [20]. In fact, it is the only TRP channel for which there is broad agreement about its central role in temperature detection, and the only well-stablished cold receptor. As such, TRPM8 is an obvious candidate to have mediated putative adaptations to cool and cold environments.

*TRPM8*, located on the short arm of human chromosome 2, harbours genetic diversity with potential functional and phenotypic consequences. A single-nucleotide polymorphism (SNP; rs10166942, C/T, chr2:234825093) upstream of the gene is strongly associated with migraine in Europeans, with the ancestral C allele being protective of migraine with and without aura [21‐24] with a relatively large effect (odds ratio 0.80-0.86 [22]). The precise molecular mechanism for this association remains unknown, although TRPM8 likely plays a role in pain perception at least with noxious cold stimuli and peripheral inflammation (reviewed in [25, 26]). Further, the channel mediates, for example, the analgesic effect of menthol in acute and inflammatory pain [27]. Genetic variation of *TRPM8* is thus likely to affect thermal sensation, which could mediate adaptations to ambient temperature. Here, we use a combination of genetic methods to resolve the evolutionary history of *TRPM8* in human populations and show strong evidence for local adaptation that correlates with latitude and temperature.

## Materials & Methods

### The rs10166942 T allele

The variant rs10166942 is located ~1 kb upstream of the *TRPM8* gene. We used a combination of bioinformatics tools to investigate possible functional effects of rs10166942 and it neighbouring variants in high linkage disequilibrium (LD). We explored the predicted effects on protein sequence using variant effect predictor (VEP) [28], focusing on the non-synonymous and splice-site SNPs, as well as indels annotated in the 1000 Genomes data (hereafter 1KGP). We explored effects on gene expression using Regulome DB annotations [29], GTEx data [30] and basal root ganglion RNA-Seq data (kindly provided by G. Gisselmann) [31].

### Modern genomes

To investigate the patterns of genetic diversity of TRPM8 we used genome-wide genotype data from the 1KGP phase III [32]. African ancestry: ESN (Esan in Nigeria), GWD (Gambian (Mandinka) in Western Divisions in Gambia), YRI (Yoruba in Ibadan, Nigeria), LWK (Luhya in Webuye, Kenya), MSL (Mende in Sierra Leone), ASW (African Ancestry in Southwest USA), ACB (African Caribbean in Barbados); European ancestry: GBR (British from England and Scotland), CEU (Utah Residents, USA, with Northern and Western European ancestry), FIN (Finnish from Finland), TSI (Toscani in Italia), IBS (Iberian Populations in Spain); East Asian ancestry: CHS (Southern Han Chinese), CHB (Han Chinese in Beijing, China), JPT (Japanese in Toyko, Japan), CDX (Chinese Dai in Xishuangbanna, China), KHV (Kinh in Ho Chi Minh City, Vietnam); South Asian ancestry: BEB (Bengali in Bangladesh), GIH (Gujarati Indians in Houston, USA), ITU (Indian Telugu in the UK), PJL (Punjabi in Lahore, Pakistan), STU (Sri Lankan Tamil in the UK). The American populations from the 1KGP have recent admixture with Europeans [33], and thus are not suited for our analysis and were excluded. Across the 22 populations the lowest sample size is 61 (ASW), so to minimise power differences among populations we randomly down-sampled each population to 61 unrelated individuals.

We also used the data from the 142 populations of the Simons Genome Diversity Panel (SGDP) project dataset, which was obtained, together with its meta-information (including geographic location) [34]. For the geographic location, in the southern hemisphere we used the absolute value of the latitude. Most populations have high coverage whole-genome sequencing data for two representative individuals, so we used two individuals from each ‘Panel C’ population with a sample size of at least two (110 populations).

### Early Eurasian genomes

Ancient genomes were used to infer the frequency of rs10166942 T in different pre-historic human populations. The genotype data from ancient paleo-eskimo individuals from the Saqqaq culture [35] were obtained from the Danish bioinformatics center. Data on early Europeans [36] was downloaded from the Reich lab webpage (see URLs). We transformed the binary eigenstrat file to a vcf using eigenstrat2vcf.py and extracted the genotype information for rs10166942. Age information was extracted from Supplementary Data 1 in [36]. After filtering, we were able to genotype 79 ancient individuals for rs10166942. These individuals lived in Eurasia 3,000 to 8,500 years ago and represent three different ancestry groups: Hunter-Gatherers (8 individuals), Early Farmers (33 individuals), and Steppe pastoralists (38 individuals).

### Origin of the rs10166942 T allele

We inferred the likely place of origin for the rs10166942 T allele by analysing haplotypes carrying the derived T allele, as levels of linked variation should be highest in the population closest to the one where it appeared. Since no homozygous T/T individuals are present in several of the 1KGP populations, we relied on the phased haplotypes across the 65 kb region of interest. We calculated pi after removing derived haplotypes with evidence of recombination with ancestral rs10166942 C allele (Table S1).

### Latitude and temperature estimates

In order to investigate the correlation of allele frequencies with latitude and temperature, we jointly analysed genetic, latitude, and temperature information. For modern humans, we estimated the absolute latitude of the location of each population according to Wikipedia and Google Maps (Table 1). The CEU population, of central European ancestry, was assigned the coordinates of Brussels. For early modern humans, latitude information was extracted from Supplementary Data 1 in [36] and updated when necessary (e.g., some individuals lacked geographic coordinates or had problems with the longitude/latitude information).

Temperature time series information was extracted for 2001-2010 from a 0.5°x0.5° grid matrix assembled at the Climate Research Unit of the University of East Anglia (version 3.23; [37]). Data is available since 1960, but we used only the time series from 2001-2010 to guarantee comparable and high-quality estimates across populations. Using the geographic coordinates of each population we extracted annual mean temperatures.

### Phylogenetic Generalized Least Squares (PGLS)

To investigate to what extent shared ancestry, latitude and temperature predict rs10166942 T allele frequency in each population we used two different linear models. We first used a Phylogenetic Generalized Least Squares (PGLS) analysis [38], which can account for the full phylogenetic signal (the population relationships) present in our data. The response variable is the mean derived allele frequency of the rs10166942 T allele per population. We first conducted a null/full model comparison. The null model contains only the shared ancestry information (the ‘phylogeny’); here, we used the full pairwise F_ST_ matrix averaged across all positions polymorphic in that particular population pair. Following Weir and Cockerham, we calculated the genome-wide average F_ST_ between two populations as the “ratio of averages” (equation 10 in [39]). A neighbor-joining (NJ) tree was calculated using a matrix of the pairwise F_ST_ values with the R package *ape* [40], and rooted using ‘mid-point’ rooting with *Archaeopteryx* [41]. The full model includes additional predictor variables: *latitude* and annual mean *temperature*. In order to achieve convergence of the model we z-transformed each predictor. We excluded populations one at a time and compared the model estimates derived from the subsets with those obtained from the full data set, which revealed the model to have good stability. We assessed for the full model whether the assumptions of normally distributed and homogenous residuals were fulfilled by visual inspection of a QQ-plot of the residuals and residuals plotted against fitted values [42], which revealed no issues with these assumptions. As an overall test of the effect of the two test predictors (*latitude* and annual mean *temperature*), we compared the fit of the full model with that of the null model [43] using a likelihood ratio test [44].

We then performed a multi-model inference [45] to compare the null model and all possible models that could be constructed with the two test predictors (four models in total). To quantify the relative performance of each model, we used Akaike’s Information Criterion (AIC, corrected for small samples) as a measure of model fit penalized for model complexity and determined Akaike weights as a measure of the support a model received compared to all other models in the set [45]. In practice, we use the Akaike weights to derive the 95% best model confidence (comprising the truly best model in the model set with a probability of 0.95) and also to determine Akaike weights for the individual predictors by summing the Akaike weights of the models comprising them. To infer the overall relevance of predictors in the model set we determined whether the null model was included in the 95% best model confidence set [46]. The analysis was conducted in R [47] using the function pgls of the package caper [48].

### Generalized Linear Mixed Models

To be able to analyze both the 1KGP and the SGDP datasets (which has low sample size for a large number of populations, so allele frequencies cannot be estimated) we also used a generalized linear mixed model [49] (GLMM) fitted with binomial error structure and logit link function [50]. This model conceptually corresponds to a regression; however, it allows more flexibility with regard to the distribution of the response (e.g., normality and homogeneity of the residuals are not necessarily required), and it also allows us to effectively control for non-independence of the data due to multiple observations of the same populations or individuals [49]. The response variable is the genotype of rs10166942 in each individual, in a 2-column-reponse-matrix (the derived and the ancestral allele counts). For the modern human genetic data, shared ancestry was controlled by adding as an additional fixed effect the genetic distance between each population and YRI, measured as the genome-wide average F_ST_. Population identity was included as a random effect in the model, to account for random genetic drift. We further included a random effect per individual to account for the non-independence of the ancestral and derived allele counts. The model that includes all these effects is the null model.

To test for the effects of *latitude* and the annual mean *temperature* we included them as test predictor variables with fixed effects. In the analysis of the early Europeans, we added *age* as a further test predictor variable. For the comparison among models (multi model inference [45]) we considered the null model and all possible models that could be constructed with the two test predictors, totalling four models (eight in the early European analysis). We assessed model stability as in case of the PGLS, which revealed the model to have good stability (Table S2). Overdispersion was no issue (dispersion parameter of the full model in the 1KGP: 0.97 and the SGDP: 0.67). The models were fitted in R [47] using the library ‘lme4’ [51].

### Signatures of local adaptation

Local adaptation on a single variant can lead to a rapid rise in the frequency of the positively selected allele, resulting in strong population differentiation (measured for example by F_ST_) between the population(s) with positive selection and those without it. We calculated per SNP F_ST_ with a custom *perl* implementation of the Weir and Cockerham estimator [39] for each pairwise population comparison.

The allele under positive selection will rise in frequency together with its background haplotype, raising the frequency of linked alleles. When the favoured allele is young (e.g., under a classic selection from a de-novo mutation model (SDN) hard sweep model), this results in a signature of extended haplotype homozygosity. To test for such signature, we calculated iHS [52] and XP-EHH [53] using *selscan* with default parameters [54]. For iHS, we used SNPs with derived allele frequencies higher than 5% and lower than 95%. For XP-EHH, we used SNPs with derived allele frequency higher than 5% in the test population. These filters follow previously established methods [55] and prevent signatures of extended LD to be broken by rare variants, while still obtaining XP-EHH values for derived alleles fixed or nearly fixed in the *test* population. For both analyses, only sites with a high confidence inferred ancestral allele were used (part of 1KGP genotype files). Recombination was estimated using the genetic map from HapMap Project, Phase 2 [56].

All three statistics were calculated genome-wide, and P-values for SNPs of interest were calculated based on the empirical distribution. Since both tests are sensitive for positive selection, the tail of the empirical distribution is enriched for the targets of positive selection. Our analysis is hypothesis-driven for the migraine risk allele in rs10166942, and, thus, no correction for multiple testing is required.

### Approximate Bayesian Computation analysis

To infer the selective history of the gene, we used an approximate bayesian computation (ABC) approach, which allows us to assess the probability of different evolutionary models and their associated parameters [57]. Following [7, 58], we compared the genomic observations to simulations under three models with parameters drawn from uniform (U) prior distributions. These models are: (I) SDN, where the selected allele appeared as a single copy between 60,000 and 30,000 years ago (ya) (t_mut_~U(30,000, 60,000ya)) and was immediately advantageous with a selective coefficient that was allowed to differ between the African (s_A_~U(0,1.5%)) and the non-African (s_NA_~U(0.5,5%)) populations; (II) selection on standing variation (SSV), where a previously neutral allele at a given starting frequency (f_sel_~U(0,20%)) became positively selected (s_NA_~U(>0,5%)) in the non-African population after the out of Africa migration and before the European-Asian split (51,000 to 21,000ya; t_mut_~U(21,000, 51,000ya)); (III) fully neutral model (NTR), where the allele appeared as in the SDN model (t_mut_~U(30,000, 60,000ya)) but was completely neutral.

We ran one million simulations for each selection model and 100,000 simulations for the neutral model using msms [59]. Each simulation comprised a stretch of 185 kb with 122 chromosomes of an African (population 1) and a non-African (population 2) population. Human demographic parameters followed the model inferred by Gravel et al. [60], and in each simulation we analyze the African population with one non-African population (in Europe or Asia). To simulate the recombination hotspots across the locus, we simulated extended regions with a length that corresponded to the local increase in recombination rate above the baseline recombination rate (Figure S1). These regions were then removed before calculating summary statistics, such that they contribute recombination events but not mutation events to the data. The baseline recombination rate was the mean recombination rate across the locus excluding the peaks, based on a merged map from several 1KGP populations (Figure S1).

For the ABC inference we used five summary statistics: XP-EHH, Fay and Wu’s H [61], Tajima’s D [62], F_ST_ [39] and derived-allele-frequency. XP-EHH and F_ST_ were calculated between YRI and the studied population. We calculated the LD based statistic XP-EHH on the selected allele using the entire simulated region. We calculated the statistics Fay and Wu’s H, Tajima’s D, and average F_ST_ (across SNPs in a section) in both simulated populations on two separate sections: the first section was the central ~65 kb part (since the genomic data shows strong population differentiation across 65 kb), and the second section were the combined flanking regions, together 120 kb long. We also used the allele frequency of the selected site in the African and non-African population and its F_ST_.

As in the genomic data, for the XP-EHH statistic we required the variant investigated to have a derived allele frequency > 5% in the *test* non-African population. The absence of a long haplotype associated with the derived allele (XP-EHH) in the presence of strong population differentiation is an important attribute to differentiate between the SDN and the SSV model [63‐65]. Thus, we used only simulations where XP-EHH could be calculated, which biased minimally the previously uniform prior.

All summary statistics were calculated in the same way for the simulations and the real data –where rs10166942 was used as a proxy for the selected site. The demographic history follows the [60] model. African demography was based on YRI, all European populations (CEU, GBR, TSI, FIN, IBS) were simulated under the inferred European (CEU) demography, and all Asian populations (CDX, CHB, CHS, KHV, JPT, BEB, GIH, ITU, PJL, STU) under the inferred East Asian (CHB/JPT) demography. The ABC analysis was performed using the ABCtoolbox on BoxCox and PLS transformed summary statistics (following recommendations for ABCtoolbox) [66] retaining the top 1,000 simulations matching our observation. We used the first five PLS components as they carried most information for each parameter (Figure S2).

The PLS transformed statistics differentiate between the different models and capture the variation observed (Figure S3), rendering them well-suited for the inference.

### Data availability statement

No datasets were generated during the current study.

### Code availability statement

Code is available upon request.

## Results

To investigate the recent evolutionary history of *TRPM8*, we focused on the rs10166942 SNP following several lines of evidence that suggest functional relevance. The first one is association with disease, as the ancestral T allele shows strong association with reduced risk of migraine [67] that has been consistently replicated in different populations e.g. [21‐23, 68]. Despite the obvious interest of these results, to date the molecular mechanism responsible for these associations remains unknown. This is most likely due to the restricted tissue expression of the gene and the temperature/ligand-dependent activation of the protein, which severely hamper experimental functional assays – as, for example, typical genome-wide experiments are run under basal conditions [69]. It is worth noting, that computational predictions suggest rs10166942 alters transcription factor binding [70]. The very specific tissue expression of the gene makes it extremely challenging to test this prediction experimentally, but a regulatory function fits well the location of the SNP, which sits ~1 kb upstream of *TRPM8*. We note that no neighbouring SNP in high linkage disequilibrium (LD) shows stronger evidence of association with migraine [22] or functionality (Figures S4) than rs10166942. Thus, rs10166942 remains as the most likely functional variant in this genomic region and we chose it as our target variant –with the understanding that we cannot discard the possibility that it tags another functional variant in this locus which would, however, share its genetic signatures.

### Latitude and *TRPM8*-rs10166942

rs10166942 shows interesting patterns of allele frequencies in the 1KGP project populations [32] (Figure 1A, Table 1). The levels of linked variation indicate that the T allele originated in Africa (Figure S6, Figure S7, Table S1) but while its frequency today is just 5% in the equatorial YRI, it reaches intermediate frequencies in Asia and up to 88% frequency in the northern European Finnish (Figure 1A, Table 1). Frequencies of the rs10166942 T allele in South Asia are on average 0.48, closer to those in East Asia (0.36) than in Europe (0.83), in contrast with the patterns of shared ancestry – genome-wide South Asian populations are closer to Europeans than to East Asians (Figure S8) [32]. Together, allele frequencies paint a seemingly latitudinal cline of allele frequencies (Figure 1A, Table 1).

**Figure 1.**
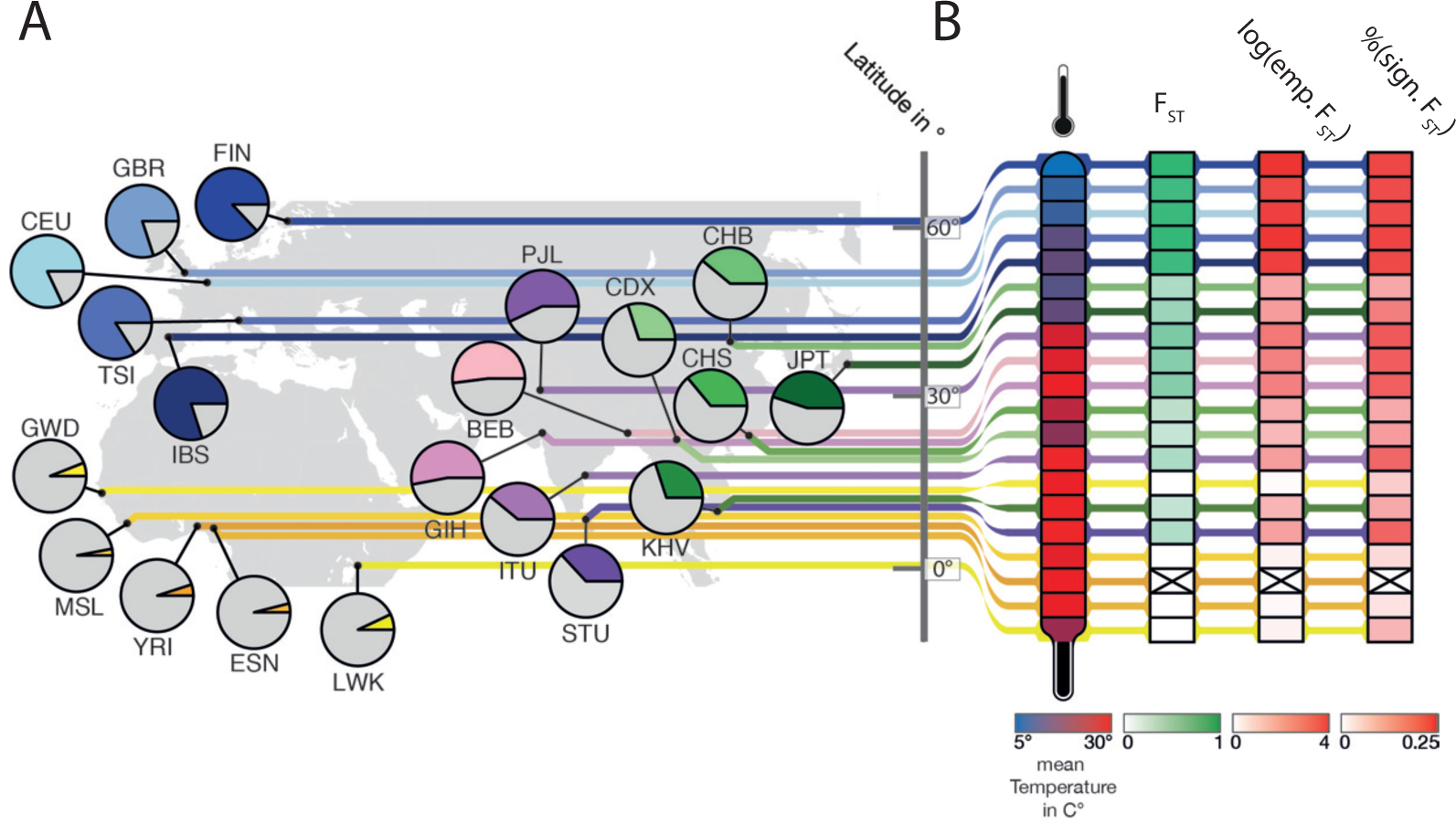
Overview of the populations used and their allele frequencies for rs10166942, average temperature, and F_ST_ signatures. **(A)** Geographic location of the 1KGP populations used, with the derived allele frequency of the rs10166942 allele in piecharts (T allele in color according to population), and their latitude. **(B)** In columns, annual mean temperature at the geographic location of each population, the level of F_ST_-based population differentiation with YRI, the log10 empirical P-value of this F_ST_ value, and the proportion of SNPs in the 65kb target region with an empirical P-value lower than 0.05.

We tested this hypothesis using linear models and, because of the thermoregulatory role of *TRPM8*, included temperature as a covariate. We tested, using a PGLS [71] analysis, to what extent shared ancestry, latitude and annual average temperature predict the observed allele frequency in each population. We first performed a model comparison between a null model (only ancestry information) and a full model (which includes latitude and temperature as predictor variables). The full model explains the data significantly better than the null model (χ^2^ = 13.04, df = 2, P-value = 0.001). When we then assessed the influence of each predictor with multi model inference, the null model again receives weak support (Table 2). The highest support is for the model with latitude, followed closely by the model with latitude and temperature; together, they make up the 95% best model confidence set (Table 2), placing latitude alone or combined with temperature as a better predictor of rs10166942 T allele frequency than shared ancestry. The correlation between allele frequency and latitude in this model is evident in Figure 2A. GLMM, which uses one-dimensional ancestry information but can use genotype data and allows non-linear fits to the data, confirmed the significant latitude correlation, with or without temperature, in 1KGP data (Figure 2B; Table 2). In addition, we confirmed this result using 110 populations of the SGDP dataset (Supplemental Data 1) [34], which provide a much denser worldwide population sample (Figure 2C, Figure S9, Table 2). Further, the significant correlation remains when only Eurasian populations are analysed (in the SGDP dataset, where the number of populations allows this analysis) showing that the inference is not driven by the low T frequencies in African populations (data not shown).

**Figure 2.**
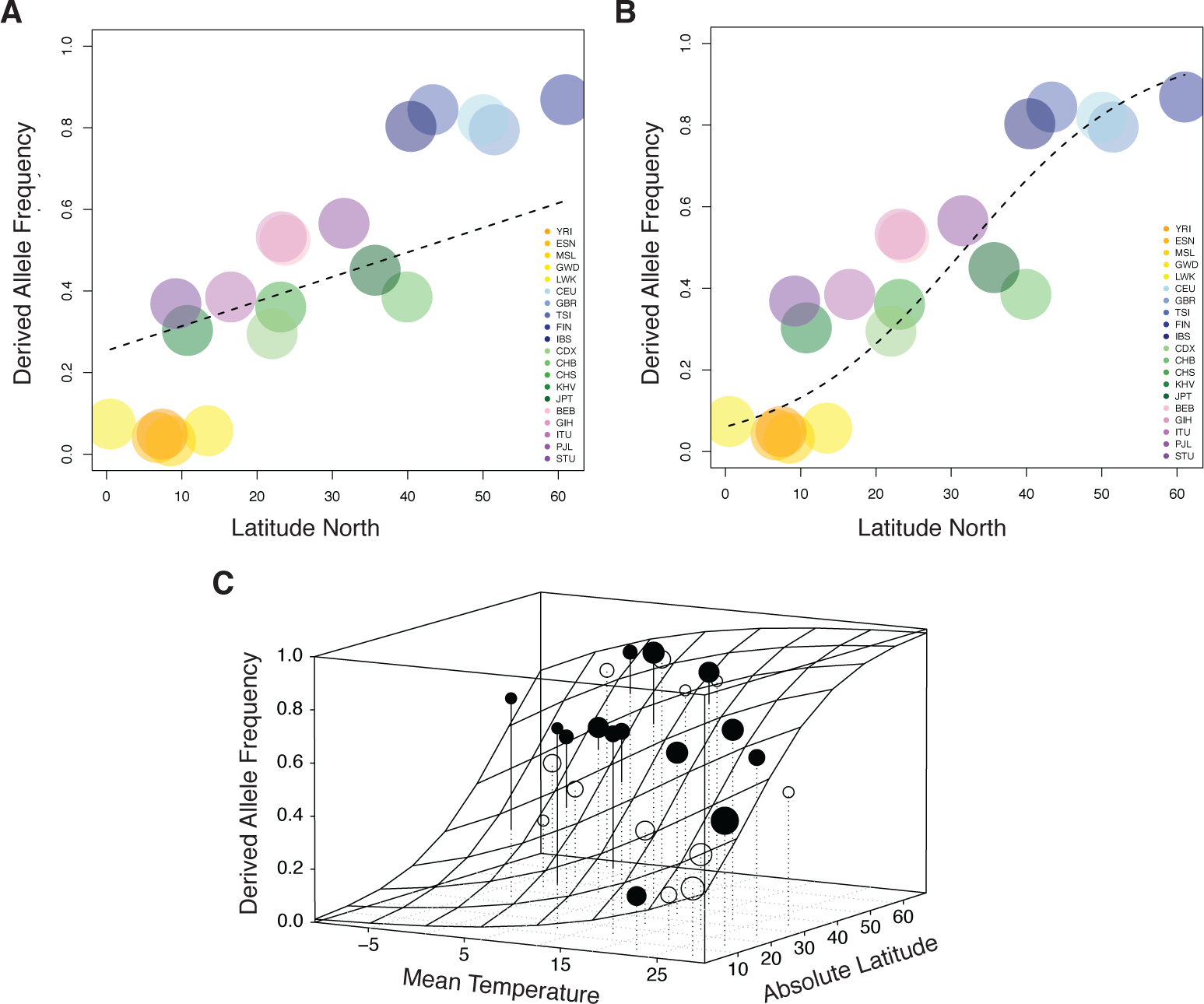
Correlation of the frequency of the rs10166942 T allele with latitude. Correlation of the frequency of the rs10166942 T allele with latitude. The fitted function (dashed line) results for the 1KG data from **(A)** the PGLS and **(B)** GLMM analysis. **(C)** Results of the best model in the PGLS analysis. The fitted response is shown as gridded surface, and the dots represent the average frequency of the rs10166942 T allele per cell of the gridded surface. Points above the surface are filled, points below are open. The volume of the points corresponds to the number of populations per cell.

Latitude is thus a strong predictor of genotype – that is, of the presence and frequency of the rs10166942 T allele in a given population. Temperature is a weaker predictor, perhaps because it is less stable over time. Available genomic data from prehistoric Eurasians (ages 3,000 to 8,500 years old [36, 72]) show no significant support for any predictor (Materials and Methods; Figure S10), although the low number and restricted geographic origin of these ancient samples markedly hamper the analysis. In any case, ancient DNA suggests that the derived rs10166942 T allele was already at appreciable frequencies in pre-historic European groups that include Hunter-Gatherers (frequency 81%), Farmers (77%), Steppe pastoralists (71%) and possibly Paleo-Eskimos from Greenland (the available genome is T homozygote) [72].

### Signatures of positive selection at *TRPM8*-rs10166942

The observation that rs10166942 frequencies are better explained by latitude than population history, with extremely high frequencies of the T allele in Northern Europe, raises the possibility that adaptation to north Eurasian environments resulted in increased frequency of this *TRPM8* allele. We first explored signatures of local positive selection using F_ST_, a measure of population differentiation to the equatorial YRI population. rs10166942 is among the most strongly differentiated SNPs genome-wide between YRI and not only all European populations (GBR, FIN, IBS, TSI, CEU; empirical P-values = 0.0002-0.0006), but also all South East Asian (STU, ITU, GIH, BEB, PJL; P-values = 0.041-0.007), and one East Asian (JPT; P-value = 0.0356) population (Figure 1B, Table 1). The high F_ST_ signature extends for ~65 kb in the upstream half of *TRPM8*, and, due to LD, some SNPs show comparable signatures, however only rs10166942 has been associated with a phenotype (Figure S11). F_ST_ sharply declines beyond the 65 kb upstream portion of *TRPM8*, probably due to recombination (Figure S11, Figure S1). Although non-African populations show relatively high LD in the locus (Figure S12), LD-based statistics show weak evidence of population-specific (XP-EHH [53]) or incomplete (iHS [52]) selective sweeps on a new advantageous mutation at rs10166942 and nearby SNPs (Table 1, Figure S11).

### Evolutionary history of TRPM8-rs10166942

The combination of unusually high F_ST_ values with ordinary LD patterns suggests that this locus evolved under recent, local positive selection but not under a classical hard selective sweep. We formally evaluated this possibility using an ABC approach, which allows us to assess the probability of different evolutionary models and their associated parameters [57]. We used the ABC, as in [7, 58], to differentiate between three models: SSV, SDN, and a neutral model (Figure 3A).

**Figure 3.**
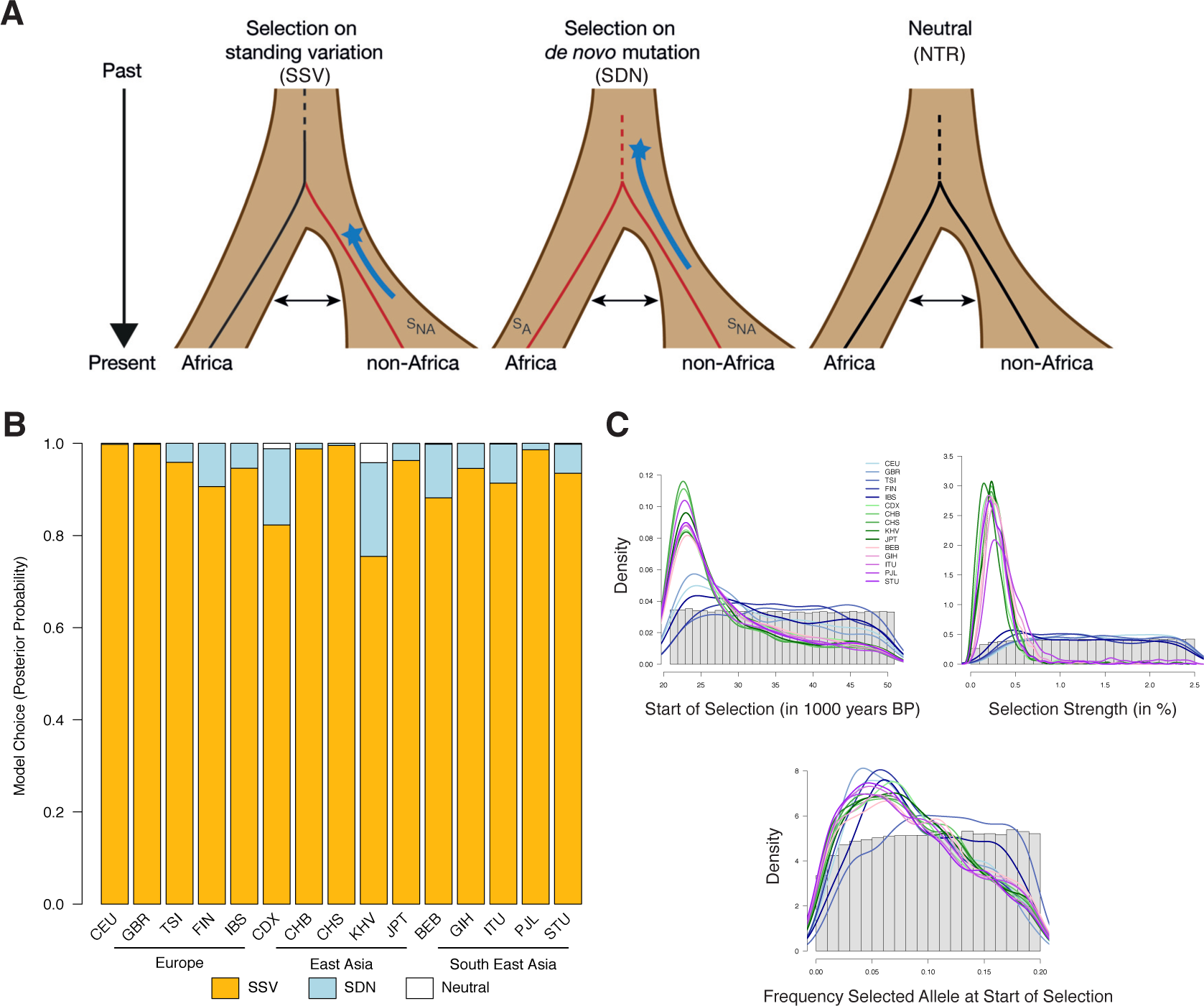
ABC analysis. Graphical representation of the three models(SSV, SDN, NTR) and their associated parameters. The range of the prior distribution for time of selection start is depicted by a star and a blue line. **(B)** Posterior probabilities for each model and population. **(C)** Prior distribution of each parameter as a histogram. Posterior distribution of the SSV model parameters as a line for each population.

We have high power to identify the correct evolutionary model (the fraction of correctly assigned simulations is 96% for SDN, 81% for SSV, and 96% for NTR) with high sensitivity and specificity (Table S3). Across all populations, the ABC results consistently favour the SSV model (Figure 3B). Bayes factors (Bayesian measure of confidence) range from 4.6 to over 500 (Table 3), representing strong to decisive evidence for the SSV model [73]. Only in KHV (2^nd^ most southern non-African population) the model choice result is inconclusive, although the SSV model still has the strongest support (Figure 3B). Interestingly, the support for the SSV model correlates moderately (almost significantly) with latitude (Pearson correlation r=0.49, p=0.06) because the signatures of selection are stronger at higher latitudes, as expected if the selective advantage of the T allele grew with latitude.

The ABC framework allows estimation of the parameters of the SSV model (Table 3), although these always have large confidence intervals so median point estimates should be taken with caution. We infer that selection started about 26,000 years ago on an allele that was at a moderate frequency (the estimate, 7.5%, is close to its current frequency in western Africa) (Table 3) and was moderately favourable in Asia (s_Non-Africa_=0.28%). In Europe, we could not confidently infer the strength of selection as this parameter’s posterior distribution is quite flat (Figure 3C). This is because selection coefficients higher than 0.5 lead to almost identical summary statistic distributions (Figure S3). However, selection strength was likely higher than 0.5 in European populations (posterior probability = 0.88), whereas in Asian populations there is little support for such high selection (posterior probability = 0.12). Together, the ABC results provide strong evidence for positive selection on neutral standing variation in all non-African populations, albeit with different selection intensities in different human groups.

## Discussion

Here we present evidence that the derived T allele of rs10166942 in *TRPM8* arose in frequency due to positive selection in a latitude-related manner. We note that while rs10166942 T is the most likely target of selection, we cannot discard that selection targeted an unknown, strongly linked allele –but this should not substantially affect our inferences. The SNP shows unusually high levels of population differentiation – it is among the 0.02% most differentiated alleles between the Yoruba and Finnish populations. Although there is a distinctive signature of high LD in the region in non-Africans, the patterns do not show clear evidence of an incomplete, hard sweep of positive selection. In fact, we infer that the derived T allele appeared in Africa and segregated neutrally, and only after the out-of-Africa migration moderate positive selection rose the standing T allele in non-African populations. ABC parameter inferences are noisy and have large confidence intervals, but our point estimates indicate that selection began about 26,000 years ago, incidentally coinciding with the last glacial maximum around 26,500 years ago [74]. According to our results, selection was moderate in Asian populations and probably stronger in Europeans. This agrees well with the high frequency of the T allele in the genomes of prehistoric Europeans.

Latitude, with or without temperature, predicts the rs10166942 allele frequency better than population history (the full phylogeny for PGLS, pairwise differentiation for GLMM) in both datasets analysed. Together with the F_ST_ signatures and ABC inferences, this suggests positive selection along a latitudinal cline raised the frequency of the rs10166942 T allele. We note, however, that even under comparable environmental pressure for one factor, alleles do not necessarily reach similar frequencies across populations, as many other environmental factors differ and contribute to the overall allele-frequency. In fact, while the latitudinal cline is significant latitude and frequency do not correlate perfectly, so additional environmental factors may be at play (perhaps in Asian populations; Figure 1, Figure S9).

Given the function of TRPM8, the cold temperatures in northern latitudes are particularly likely to drive positive selection in this locus. The fact that overall current average temperature is a weaker predictor of allele frequency than latitude could be due to the considerable fluctuations of temperature over time (here, thousands of years) and the fact that the recorded data (monthly averages) is not particularly informative about long-term selective pressures. Latitude is strongly correlated with numerous other aspects of climate and is likely a good proxy for the long-term effects of climate in each of the human populations analysed, perhaps even better than current temperature. Nevertheless, it remains possible that other unknown functions of TRPM8 have mediated the allele frequency change. For instance, a gastrointestinal role has been described for TRPM8 [75] as well as an association of rs10166942 with inflammatory bowel syndrome [70], however, its expression in the gut has not been unequivocally established [76].

As mentioned above, rs10166942 is also among the most strongly associated SNPs with migraine incidence genome-wide [21‐23]. Migraine is a debilitating neurological disorder that affects millions of people worldwide [77]. While several non-genetic traits increase the individual risk of migraine, notably being of middle age, female, suffering high stress levels and having a low socio-economic status [78, 79], genetics play an important role. In fact, migraine is a highly heritable (34% - 57% heritability [80]) yet polygenic disease [23]. Given the association between rs10166942 C and low risk of migraine, the adaptive local rise in frequency of the T allele (due to direct positive selection or linkage to a selected site) could have contributed, to some extent, to differences in migraine prevalence in certain human groups. This agrees with epidemiological data: according to the World Health Organization, migraine shows low prevalence in Africa, highest prevalence in Europe, and intermediate prevalence in the Asian countries at intermediate latitudes among the two [77, 81]. In the USA migraine prevalence has consistently been shown to be higher for European-Americans than African-Americans after non-genetic confounding factors are accounted for [81, 82]. Although the putative influence of rs10166942 in migraine risk is moderate, and additional factors are likely at play, migraine prevalence correlates with the evidence of positive selection and the frequency of the T allele. Thus, local adaptation in *TRPM8* may have contributed to modify, by yet unknown molecular mechanisms, pain-related phenotypes in human populations.

## Author Contributions

AMA conceived and supervised the study. FMK, AMA designed the experiments. MDA contributed data and information. FMK, MAA, JS explored the signatures of natural selection. RM, FMK, JS performed GLMM and PGLS analysis. MYD, FMK performed inference of the functional consequences of alleles. FMK, BP performed ABC analysis. FMK, AMA interpreted the results together with all other authors. FMK, AMA wrote the manuscript, with contributions from all authors.

## Acknowledgements

We thank E. Huerta-Sanchez, F. Romagné, M. Dannemann, I. Mathieson, and G. Gisselmann for sharing data and/or scripts. Wulf Hevers and Robert Kraft for discussing functional implications of non-synonymous SNPs. Mark Stoneking, Sergi Castellano, David Reher, and Monty Slatkin for critical comments on the manuscript. This work was funded by the Max Planck Society. MDA was supported by a grant from the Department of Health of the Basque Government (2015111133). BMP was supported through NIH R01 HG007089 and an early postdoc mobility fellowship from the Swiss NSF. MYD is supported in part through NIH R00NS083627 from the National Institute of Neurological Disorder and Stroke.

## URLs

1000Genomes genotypes

http://ftp.1000genomes.ebi.ac.uk/vol1/ftp/release/20130502

1000Genomes recombination rate

ftp://ftp.1000genomes.ebi.ac.uk/vol1/ftp/technical/working/20130507_omni_re_combination_rates/

SGDP v3

http://sharehost.hms.harvard.edu/genetics/reich_lab/cteam_lite_public3.tar

Temperature Time Series

http://browse.ceda.ac.uk/browse/badc/cru/data/cru_ts/cru_ts_3.23/data/tmp

Ancient Paleo-Eskimo

http://www.binf.ku.dk/Saqqaq

Ancient Eurasian analysis:

https://genetics.med.harvard.edu/reich/Reich_Lab/Datasets.html;

https://github.com/mathii/

## Supplemental Data Legend

**Supplemental Data 1. Overview SGDP data.** For each individual used from the SGDP ‘C Panel’ the ID, population, continent, rs10166942 ancestral and derived allele count, latitude, longitude and mean year wise temperature are given (txt file).

**Table 1.** Overview of populations and signatures of natural selection. Geographic coordinates (in degrees), mean annual temperature (in degrees Celsius), and the frequency and signatures of selection for the rs10166942 T allele (empirical P-value), per population, ordered by latitude. DAF: derived allele frequency. Continents: (EUR) Europe, (EAS) East Asia, (SEA) South East Asia, (AFR) Africa. Not calculated because YRI was used as background population. XP-EHH not calculated within Africa. Allele frequency did not meet criteria (see Methods).

| Population | Continent | Latitude | Longitude | Temperature | DAF | F<sub>ST</sub> | F<sub>ST</sub> P-value | XP-EHH P value | iHS P-value |
| --- | --- | --- | --- | --- | --- | --- | --- | --- | --- |
| FIN | EUR | 60,25N | 24,75E | 5.7 | 0.87 | 0.805 | 0.0002 | 0.205 | 0.166 |
| GBR | EUR | 54,75N | 1,25W | 10.0 | 0.80 | 0.724 | 0.0006 | 0.287 | 0.304 |
| CEU | EUR | 50,75N | 4,25E | 10.7 | 0.82 | 0.751 | 0.0004 | 0.228 | 0.36 |
| TSI | EUR | 43,25N | 11,25E | 14.2 | 0.84 | 0.778 | 0.0002 | 0.26 | 0.622 |
| IBS | EUR | 40,25N | 3,25W | 14.9 | 0.80 | 0.733 | 0.0004 | 0.291 | 0.656 |
| CHB | EAS | 39,75N | 116,25E | 13.4 | 0.39 | 0.279 | 0.0550 | 0.939 | 0.219 |
| JPT | EAS | 35,25N | 139,25E | 14.8 | 0.45 | 0.349 | 0.0356 | 0.947 | 0.593 |
| PJL | SEA | 31,25N | 74,25E | 25.3 | 0.57 | 0.472 | 0.0066 | 0.651 | 0.869 |
| BEB | SEA | 23,25N | 90,25E | 26.1 | 0.52 | 0.428 | 0.0102 | 0.605 | 0.8 |
| GIH | SEA | 23,25N | 72,75E | 27.7 | 0.53 | 0.437 | 0.0101 | 0.587 | 0.821 |
| CHS | EAS | 22,25N | 114,25E | 23.4 | 0.36 | 0.254 | 0.0666 | 0.927 | 0.161 |
| CDX | EAS | 22,25N | 100,25E | 19.2 | 0.30 | 0.184 | 0.1051 | 0.895 | 0.926 |
| ITU | SEA | 16,75N | 80,75E | 28.6 | 0.39 | 0.278 | 0.0367 | 0.804 | 0.952 |
| GWD | AFR | 13,25N | 16,25W | 27.2 | 0.06 | -0.007 | 0.8610 | NA<sup>b</sup> | 0.39 |
| KHV | EAS | 10,25N | 106,25E | 28.2 | 0.30 | 0.193 | 0.0953 | 0.938 | 0.69 |
| ESN | AFR | 6,75N | 6,25E | 27.0 | 0.04 | -0.007 | 0.8046 | NA<sup>b</sup> | NA<sup>c</sup> |
| STU | SEA | 9,25N | 80,25E | 28.5 | 0.37 | 0.262 | 0.0411 | 0.866 | 0.626 |
| MSL | AFR | 7,75N | 11,25W | 26.6 | 0.03 | -0.005 | 0.6836 | NA<sup>b</sup> | NA<sup>c</sup> |
| YRI | AFR | 7,25N | 3,75E | 27.6 | 0.05 | NA<sup>a</sup> | NA<sup>a</sup> | NA<sup>a, b</sup> | 0.699 |
| LWK | AFR | 0,75N | 34,75E | 20.5 | 0.07 | -0.002 | 0.6912 | NA<sup>b</sup> | 0.901 |

**Table 2.** PGLS and GLMM analysis. All models considered, ordered by their fit Model rank). Three measures of model support are shown: AIC, delta AIC, and Akaike weight. The cumulative Akaike weights are shown together with the cumulative probability, and resulting confidence set (models that together provide just over 0.95 cumulative Probability; indicated by ‘yes’). k: number of estimated parameters. Results using the 1000 Genomes data are shown for the PGLS, and for the GLMM, and the GLMM results for the SGDP data.

|  | Models | Model Rank | AIC | delta AIC | weight AIC | cumulative Pr. | confid. Set | k |
| --- | --- | --- | --- | --- | --- | --- | --- | --- |
| **1KGP PGLS** | Null+Lat. | 1 | -49.43 | 0 | 0.504 | 0.504 | yes | 5 |
|  | Null+Temp.+Lat. | 2 | -49.186 | 0.244 | 0.446 | 0.95 | yes | 6 |
|  | Null+Temp. | 3 | -44.147 | 5.283 | 0.036 | 0.986 | no | 5 |
|  | Null | 4 | -42.255 | 7.175 | 0.014 | 1 | no | 4 |
| **1KGP GLMM** | Null+Lat. | 1 | 1929.7 | 0 | 0.5165 | 0.5165 | yes | 5 |
|  | Null+Temp.+Lat. | 2 | 1929.8 | 0.1458 | 0.4802 | 0.9966 | yes | 6 |
|  | Null+Temp. | 3 | 1939.8 | 10.1411 | 0.0032 | 0.9999 | no | 5 |
|  | Null | 4 | 1946.2 | 16.4724 | 0.0001 | 1 | no | 4 |
| **SGDP GLMM** | Null+Temp.+Lat. | 1 | 435.206 | 0 | 0.943 | 0.943 | yes | 6 |
|  | Null+Lat. | 2 | 440.841 | 5.635 | 0.056 | 1 | yes | 5 |
|  | Null+Temp. | 3 | 451.699 | 16.494 | 0 | 1 | no | 5 |
|  | Null | 4 | 452.458 | 17.252 | 0 | 1 | no | 4 |

**Table 3.** ABC results of the SSV model for each population. Bayes factor (measure of confidence) and the resulting posterior probability (Post. Prob.) for the SSV model in each population, ordered by latitude. t_0_: time when selection starts; S_NA_: selection strength in non-African population; f_sel_: frequency of allele at selection start. The median of the posterior distribution of each inferred parameter is shown together with its 95% confidence interval (2.5% - 97.5%).

| Population | Bayes Factor | Post. Prob. | $t_0$ (in years) | | | $S_{NA}$ (in %) | | | $f_{sel}$ (in %) | | |
| --- | --- | --- | --- | --- | --- | --- | --- | --- | --- | --- | --- |
|  |  |  | median | 2.5% | 97.5% | median | 2.5% | 97.5% | median | 2.5% | 97.5% |
| FIN | 9.6 | 0.906 | 35055 | 22052 | 49881 | 1.238 | 0.304 | 2.430 | 0.078 | 0.010 | 0.189 |
| GBR | 588.3 | 0.998 | 29783 | 21384 | 49231 | 1.352 | 0.333 | 2.456 | 0.075 | 0.012 | 0.182 |
| CEU | 474.8 | 0.998 | 31390 | 21311 | 49593 | 1.453 | 0.346 | 2.425 | 0.080 | 0.012 | 0.187 |
| TSI | 23.3 | 0.959 | 36789 | 22088 | 50000 | 1.418 | 0.304 | 2.446 | 0.111 | 0.018 | 0.194 |
| IBS | 17.5 | 0.946 | 32558 | 21520 | 49666 | 1.209 | 0.250 | 2.409 | 0.090 | 0.012 | 0.191 |
| CHB | 81.8 | 0.988 | 24,529 | 21067 | 47771 | 0.270 | 0.045 | 0.693 | 0.081 | 0.008 | 0.191 |
| JPT | 25.8 | 0.963 | 25509 | 21103 | 48685 | 0.269 | 0.050 | 0.734 | 0.080 | 0.008 | 0.193 |
| PJL | 70 | 0.986 | 25017 | 21101 | 48055 | 0.378 | 0.109 | 2.100 | 0.077 | 0.006 | 0.192 |
| BEB | 7.4 | 0.882 | 26887 | 21118 | 48393 | 0.293 | 0.075 | 0.713 | 0.082 | 0.006 | 0.193 |
| GIH | 17.4 | 0.946 | 26,298 | 21122 | 48234 | 0.314 | 0.087 | 0.836 | 0.079 | 0.006 | 0.192 |
| CHS | 220.9 | 0.996 | 24407 | 21047 | 47586 | 0.271 | 0.049 | 0.711 | 0.079 | 0.007 | 0.189 |
| CDX | 4.6 | 0.823 | 26438 | 21088 | 47862 | 0.234 | 0.033 | 1.103 | 0.075 | 0.005 | 0.187 |
| ITU | 10.6 | 0.914 | 26,297 | 21100 | 48424 | 0.249 | 0.041 | 0.991 | 0.073 | 0.005 | 0.189 |
| KHV | 3.1 | 0.755 | 26399 | 21110 | 48452 | 0.204 | 0.025 | 1.041 | 0.075 | 0.005 | 0.186 |
| ITU | 10.6 | 0.914 | 26,297 | 21100 | 48424 | 0.249 | 0.041 | 0.991 | 0.073 | 0.005 | 0.189 |

## Bibliography

Cardona A, Pagani L, Antao T, Lawson DJ, Eichstaedt CA, Yngvadottir B, et al. Genome-wide analysis of cold adaptation in indigenous Siberian populations. PLoS One. 2014;9(5):e98076. doi: 10.1371/journal.pone.0098076. PubMed PMID: 24847810; PubMed Central PMCID: PMCPMC4029955.

Clemente FJ, Cardona A, Inchley CE, Peter BM, Jacobs G, Pagani L, et al. A Selective Sweep on a Deleterious Mutation in <i> CPT1A</i> in Arctic Populations. The American Journal of Human Genetics. 2014.

Fumagalli M, Moltke I, Grarup N, Racimo F, Bjerregaard P, Jorgensen ME, et al. Greenlandic Inuit show genetic signatures of diet and climate adaptation. Science. 2015;349(6254):1343–7. doi: 10.1126/science.aab2319. PubMed PMID: 26383953.

Racimo F, Gokhman D, Fumagalli M, Ko A, Hansen T, Moltke I, et al. Archaic adaptive introgression in TBX15/WARS2. Mol Biol Evol. 2016. Epub 2016/12/23. doi: 10.1093/molbev/msw283. PubMed PMID: 28007980.

Key FM, Fu Q, Romagné F, Lachmann M, Andrés AM. Human adaptation and population differentiation in the light of ancient genomes. Nat Commun. 2016;7:10775. doi: 10.1038/ncomms10775. PubMed PMID: 26988143.

Hancock AM, Witonsky DB, Ehler E, Alkorta-Aranburu G, Beall C, Gebremedhin A, et al. Human adaptations to diet, subsistence, and ecoregion are due to subtle shifts in allele frequency. Proceedings of the National Academy of Sciences. 2010;107(Supplement 2):8924–30. doi: 10.1073/pnas.0914625107.

Key FM, Peter B, Dennis MY, Huerta-Sánchez E, Tang W, Prokunina-Olsson L, et al. Selection on a Variant Associated with Improved Viral Clearance Drives Local, Adaptive Pseudogenization of Interferon Lambda 4 (IFNL4). PLoS Genet. 2014;10(10):e1004681. doi: 10.1371/journal.pgen.1004681.

Fumagalli M, Sironi M, Pozzoli U, Ferrer-Admettla A, Pattini L, Nielsen R. Signatures of environmental genetic adaptation pinpoint pathogens as the main selective pressure through human evolution. PLoS genetics. 2011;7(11):e1002355.

Raj SM, Pagani L, Gallego Romero I, Kivisild T, Amos W. A general linear model-based approach for inferring selection to climate. BMC Genetics. 2013;14:87. doi: 10.1186/1471-2156-14-87. PubMed PMID: 24053227; PubMed Central PMCID: PMCPMC3853933.

Wang H, Siemens J. TRP ion channels in thermosensation, thermoregulation and metabolism. Temperature. 2015;2(2):178–87.

Bautista DM, Siemens J, Glazer JM, Tsuruda PR, Basbaum AI, Stucky CL, et al. The menthol receptor TRPM8 is the principal detector of environmental cold. Nature. 2007;448(7150):204–8.

Colburn RW, Lubin ML, Stone DJ, Jr., Wang Y, Lawrence D, D’Andrea MR, et al. Attenuated cold sensitivity in TRPM8 null mice. Neuron. 2007;54(3):379–86. Epub 2007/05/08. doi: 10.1016/j.neuron.2007.04.017. PubMed PMID: 17481392.

Dhaka A, Murray AN, Mathur J, Earley TJ, Petrus MJ, Patapoutian A. TRPM8 is required for cold sensation in mice. Neuron. 2007;54(3):371–8.

Milenkovic N, Zhao W-J, Walcher J, Albert T, Siemens J, Lewin GR, et al. A somatosensory circuit for cooling perception in mice. Nature neuroscience. 2014;17(11):1560–6.

Peier AM, Moqrich A, Hergarden AC, Reeve AJ, Andersson DA, Story GM, et al. A TRP channel that senses cold stimuli and menthol. Cell. 2002;108(5):705–15.

Voets T, Droogmans G, Wissenbach U, Janssens A, Flockerzi V, Nilius B. The principle of temperature-dependent gating in cold-and heat-sensitive TRP channels. Nature. 2004;430(7001):748–54.

McKemy DD, Neuhausser WM, Julius D. Identification of a cold receptor reveals a general role for TRP channels in thermosensation. Nature. 2002;416(6876):52–8.

Dussor G, Yan J, Xie JY, Ossipov MH, Dodick DW, Porreca F. Targeting TRP channels for novel migraine therapeutics. ACS chemical neuroscience. 2014;5(11):1085–96.

Janssens A, Gees M, Toth BI, Ghosh D, Mulier M, Vennekens R, et al. Definition of two agonist types at the mammalian cold-activated channel TRPM8. Elife. 2016;5:e17240.

Ferrandiz-Huertas C, Mathivanan S, Wolf CJ, Devesa I, Ferrer-Montiel A. Trafficking of thermotrp channels. Membranes. 2014;4(3):525–64.

Anttila V, Winsvold BS, Gormley P, Kurth T, Bettella F, McMahon G, et al. Genome-wide meta-analysis identifies new susceptibility loci for migraine. Nature Genetics. 2013;45(8):912–7. doi: 10.1038/ng.2676.

Chasman DI, Schürks M, Anttila V, de Vries B, Schminke U, Launer LJ, et al. Genome-wide association study reveals three susceptibility loci for common migraine in the general population. Nature Genetics. 2011;43(7):695–8. doi: 10.1038/ng.856.

Gormley P, Anttila V, Winsvold BS, Palta P, Esko T, Pers TH, et al. Meta-analysis of 375,000 individuals identifies 38 susceptibility loci for migraine. Nature Genetics. 2016. doi: 10.1038/ng.3598.

Freilinger T, Anttila V, de Vries B, Malik R, Kallela M, Terwindt GM, et al. Genome-wide association analysis identifies susceptibility loci for migraine without aura. Nature genetics. 2012;44(7):777–82.

Julius D. TRP channels and pain. Annual review of cell and developmental biology. 2013;29:355–84.

Dai Y. TRPs and pain. Seminars in Immunopathology. 2016;38(3):277–91. doi: 10.1007/s00281-015-0526-0.

Liu B, Fan L, Balakrishna S, Sui A, Morris JB, Jordt S-E. TRPM8 is the principal mediator of menthol-induced analgesia of acute and inflammatory pain. PAIN®. 2013;154(10):2169–77.

McLaren W, Gil L, Hunt SE, Riat HS, Ritchie GRS, Thormann A, et al. The Ensembl Variant Effect Predictor. Genome Biology. 2016;17(1):122. doi: 10.1186/s13059-016-0974-4.

Boyle AP, Hong EL, Hariharan M, Cheng Y, Schaub MA, Kasowski M, et al. Annotation of functional variation in personal genomes using RegulomeDB. Genome Research. 2012;22(9):1790–7. doi: 10.1101/gr.137323.112. PubMed PMID: 22955989.

Consortium G. The Genotype-Tissue Expression (GTEx) pilot analysis: Multitissue gene regulation in humans. Science. 2015;348(6235):648–60.

Flegel C, Schöbel N, Altmüller J, Becker C, Tannapfel A, Hatt H, et al. RNA-Seq Analysis of Human Trigeminal and Dorsal Root Ganglia with a Focus on Chemoreceptors. PLOS ONE. 2015;10(6):e0128951. doi: 10.1371/journal.pone.0128951.

Consortium GP. A global reference for human genetic variation. Nature. 2015;526(7571):68–74.

Gravel S, Zakharia F, Moreno-Estrada A, Byrnes JK, Muzzio M, Rodriguez-Flores JL, et al. Reconstructing Native American migrations from whole-genome and whole-exome data. PLoS Genet. 2013;9(12):e1004023.

Mallick S, Li H, Lipson M, Mathieson I, Gymrek M, Racimo F, et al. The Simons Genome Diversity Project: 300 genomes from 142 diverse populations. Nature. 2016;538(7624):201–6. doi: 10.1038/nature18964 http://www.nature.com/nature/journal/v538/n7624/abs/nature18964.html-supplementary-information.

Rasmussen M, Li Y, Lindgreen S, Pedersen JS, Albrechtsen A, Moltke I, et al. Ancient human genome sequence of an extinct Palaeo-Eskimo. Nature. 2010;463. doi: 10.1038/nature08835.

Mathieson I, Lazaridis I, Rohland N, Mallick S, Patterson N, Roodenberg SA, et al. Genome-wide patterns of selection in 230 ancient Eurasians. Nature. 2015;528(7583):499–503.

High Resolution Gridded Data of Month-by-month Variation in Climate (Jan. 1901- Dec. 2014). [Internet]. 2015.

Grafen A. The phylogenetic regression. Philosophical Transactions of the Royal Society of London Series B, Biological Sciences. 1989;326(1233):119–57.

Weir BS, Cockerham CC. Estimating F-Statistics for the Analysis of Population Structure. Evolution. 1984;38(6):1358. doi: 10.2307/2408641.

Paradis E, Claude J, Strimmer K. APE: analyses of phylogenetics and evolution in R language. Bioinformatics. 2004;20(2):289–90.

Han MV, Zmasek CM. phyloXML: XML for evolutionary biology and comparative genomics. BMC Bioinformatics. 2009;10:356-. doi: 10.1186/1471-2105-10-356. PubMed PMID: PMC2774328.

Mundry R. Statistical issues and assumptions of phylogenetic generalized least squares. Modern phylogenetic comparative methods and their application in evolutionary biology: Springer; 2014. p. 131–53.

Forstmeier W, Schielzeth H. Cryptic multiple hypotheses testing in linear models: overestimated effect sizes and the winner’s curse. Behavioral Ecology and Sociobiology. 2011;65(1):47–55. doi: 10.1007/s00265-010-1038-5. PubMed PMID: WOS:000285786000005.

Dobson AJ, Barnett AG. An Introduction to Generalized Linear Models Third Edition Introduction. Ch Crc Text Stat Sci. 2008;77:1-+. PubMed PMID: WOS:000266971200001.

Burnham KP, Anderson DR. Multimodel inference - understanding AIC and BIC in model selection. Sociol Method Res. 2004;33(2):261–304. doi: 10.1177/0049124104268644. PubMed PMID: WOS:000224706300004.

Mundry R. Issues in information theory-based statistical inference-a commentary from a frequentist’s perspective. Behavioral Ecology and Sociobiology. 2011;65(1):57–68. doi: 10.1007/s00265-010-1040-y. PubMed PMID: WOS:000285786000006.

Team RC. R: A Language and Environment for Statistical Computing. 2016.

Orme D. The caper package: comparative analysis of phylogenetics and evolution in R. R package version. 2013;5(2).

Baayen RH. Analyzing linguistic data : a practical introduction to statistics using R. Cambridge, UK; New York: Cambridge University Press; 2008. xiii, 353 p. p.

McCullagh P, Nelder JA. Generalized linear models. 2nd ed. Boca Raton: Chapman & Hall/CRC; 1998. xix, 511 p. p.

Douglas Bates MM, Ben Bolker, Steve Walker. Fitting Linear Mixed-Effects Models Using lme4. Journal of Statistical Software. 2015;67(1):1–48. doi: doi: 10.18637/jss.v067.i01.

Voight BF, Kudaravalli S, Xiaoquan W, Pritchard JK. A Map of Recent Positive Selection in the Human Genome. PLoS Biol. 2006;4(3):e72. doi: 10.1371/journal.pbio.0040072.

Sabeti PC, Varilly P, Fry B, Lohmueller J, Hostetter E, Cotsapas C, et al. Genome-wide detection and characterization of positive selection in human populations. Nature. 2007;449(7164):913–8. doi: 10.1038/nature06250.

Szpiech ZA, Hernandez RD. selscan: an efficient multithreaded program to perform EHH-based scans for positive selection. Mol Biol Evol. 2014;31(10):2824–7. doi: 10.1093/molbev/msu211. PubMed PMID: 25015648; PubMed Central PMCID: PMCPMC4166924.

Grossman Sharon R, Andersen Kristian G, Shlyakhter I, Tabrizi S, Winnicki S, Yen A, et al. Identifying Recent Adaptations in Large-Scale Genomic Data. Cell. 2013;152(4):703–13. doi: 10.1016/j.cell.2013.01.035.

Frazer KA, Ballinger DG, Cox DR, Hinds DA, Stuve LL, Gibbs RA, et al. A second generation human haplotype map of over 3.1 million SNPs. Nature. 2007;449(7164):851–61. doi: 10.1038/nature06258.

Beaumont MA, Zhang W, Balding DJ. Approximate Bayesian computation in population genetics. Genetics. 2002;162(4):2025–35.

Peter B, Huerta-Sanchez E, Nielsen R. Distinguishing between Selective Sweeps from Standing Variation and from a De Novo Mutation. PLoS Genet. 2012;8(10):e1003011. doi: 10.1371/journal.pgen.1003011.

Ewing G, Hermisson J. MSMS: a coalescent simulation program including recombination, demographic structure and selection at a single locus. Bioinformatics. 2010;26(16):2064–5.

Gravel S, Henn BM, Gutenkunst RN, Indap AR, Marth GT, Clark AG, et al. Demographic history and rare allele sharing among human populations. Proceedings of the National Academy of Sciences. 2011;108(29):11983–8. doi: 10.1073/pnas.1019276108.

Fay JC, Wu CI. Hitchhiking under positive Darwinian selection. Genetics. 2000;155(3):1405.

Tajima F. Statistical method for testing the neutral mutation hypothesis by DNA polymorphism. Genetics. 1989;123(3):585.

Przeworski M, Coop G, Wall JD. The Signature of positive selection on standing genetic variation. Evolution. 2005;59(11):2312–23. doi: 10.1111/j.0014-3820.2005.tb00941.x.

Sabeti PC, Schaffner SF, B. Fry, J. Lohmueller, P. Varilly, O. Shamovsky, et al. Positive Natural Selection in the Human Lineage. Science. 2006 June 6:1614.

Hermisson J, Pennings PS. Soft sweeps molecular population genetics of adaptation from standing genetic variation. Genetics. 2005;169(4):2335–52.

Wegmann D, Leuenberger C, Neuenschwander S, Excoffier L. Abctoolbox: a versatile toolkit for approximate bayesian computations. BMC bioinformatics. 2010;11(1):116.

Anttila V, Stefansson H, Kallela M, Todt U, Terwindt GM, Calafato MS, et al. Genome-wide association study of migraine implicates a common susceptibility variant on 8q22.1. Nature Genetics. 2010;42(10):869–73. doi: 10.1038/ng.652.

Esserlind AL, Christensen AF, Le H, Kirchmann M, Hauge AW, Toyserkani NM, et al. Replication and meta-analysis of common variants identifies a genome-wide significant locus in migraine. European journal of neurology. 2013;20(5):765–72. Epub 2013/01/09. doi: 10.1111/ene.12055. PubMed PMID: 23294458.

Dhaka A, Viswanath V, Patapoutian A. Trp ion channels and temperature sensation. Annu Rev Neurosci. 2006;29:135–61.

Henstrom M, Hadizadeh F, Beyder A, Bonfiglio F, Zheng T, Assadi G, et al. TRPM8 polymorphisms associated with increased risk of IBS-C and IBS-M. Gut. 2016. Epub 2016/12/16. doi: 10.1136/gutjnl-2016-313346. PubMed PMID: 27974553.

Freckleton RP, Harvey PH, Pagel M. Phylogenetic analysis and comparative data: a test and review of evidence. The American Naturalist. 2002;160(6):712–26.

Rasmussen M, Li Y, Lindgreen S, Pedersen JS, Albrechtsen A, Moltke I, et al. Ancient human genome sequence of an extinct Palaeo-Eskimo. Nature. 2010;463(7282):757–62.

Jeffreys H. The theory of probability: Oxford University Press; 1998.

Clark PU, Dyke AS, Shakun JD, Carlson AE, Clark J, Wohlfarth B, et al. The Last Glacial Maximum. Science. 2009;325(5941):710–4. doi: 10.1126/science.1172873.

Zhang L, Jones S, Brody K, Costa M, Brookes SJ. Thermosensitive transient receptor potential channels in vagal afferent neurons of the mouse. American journal of physiology Gastrointestinal and liver physiology. 2004;286(6):G983–91. Epub 2004/01/17. doi: 10.1152/ajpgi.00441.2003. PubMed PMID: 14726308.

Penuelas A, Tashima K, Tsuchiya S, Matsumoto K, Nakamura T, Horie S, et al. Contractile effect of TRPA1 receptor agonists in the isolated mouse intestine. European journal of pharmacology. 2007;576(1–3):143–50. Epub 2007/09/11. doi: 10.1016/j.ejphar.2007.08.015. PubMed PMID: 17825279.

Organization WH. Atlas of headache disorders and resources in the world 2011: World Health Organisation; 2011.

Stewart WF, Lipton RB, Celentano DD, Reed ML. Prevalence of Migraine Headache in the United-States - Relation to Age, Income, Race, and Other Sociodemographic Factors. Jama-J Am Med Assoc. 1992;267(1):64–9. doi: DOI 10.1001/jama.267.1.64. PubMed PMID: WOS:A1992GW84800025.

Stewart WF, Simon D, Shechter A, Lipton RB. Population variation in migraine prevalence: a meta-analysis. Journal of clinical epidemiology. 1995;48(2):269–80.

Mulder EJ, Van Baal C, Gaist D, Kallela M, Kaprio J, Svensson DA, et al. Genetic and environmental influences on migraine: a twin study across six countries. Twin Res. 2003;6(5):422–31. doi: 10.1375/136905203770326420. PubMed PMID: 14624726.

Stovner L, Hagen K, Jensen R, Katsarava Z, Lipton R, Scher A, et al. The global burden of headache: a documentation of headache prevalence and disability worldwide. Cephalalgia. 2007;27(3):193–210. doi: 10.1111/j.1468-2982.2007.01288.x.

Stewart WF, Lipton RB, Liberman J. Variation in migraine prevalence by race. Neurology. 1996;47(1):52–9. doi: 10.1212/WNL.47.1.52. PubMed PMID: 8710124.

